# OLGenie: Estimating Natural Selection to Predict Functional Overlapping Genes

**DOI:** 10.1101/2019.12.14.876607

**Authors:** Chase W. Nelson, Zachary Ardern, Xinzhu Wei

## Abstract

Purifying (negative) natural selection is a hallmark of functional biological sequences, and can be detected in protein-coding genes using the ratio of nonsynonymous to synonymous substitutions per site (*d*_N_/*d*_S_). However, when two genes overlap the same nucleotide sites in different frames, synonymous changes in one gene may be nonsynonymous in the other, perturbing *d*_N_/*d*_S_. Thus, scalable methods are needed to estimate functional constraint specifically for overlapping genes (OLGs). We propose OLGenie, which implements a modification of the Wei-Zhang method. Assessment with simulations and controls from viral genomes (58 OLGs and 176 non-OLGs) demonstrates low false positive rates and good discriminatory ability in differentiating true OLGs from non-OLGs. We also apply OLGenie to the unresolved case of HIV-1’s putative *antisense protein* gene, showing significant purifying selection. OLGenie can be used to study known OLGs and to predict new OLGs in genome annotation. Software and example data are freely available at https://github.com/chasewnelson/OLGenie.

## Introduction

Natural selection in protein-coding genes is commonly inferred by comparing the number of nonsynonymous (amino acid changing; *d*_N_) and synonymous (not amino acid changing; *d*_S_) substitutions per site, with *d*_N_/*d*_S_ < 1 indicative of purifying (negative) selection. Thus, *d*_N_/*d*_S_ can be used to predict functional genes (Gojobori et al. 1982; Nekrutenko et al. 2002). However, complications arise if synonymous changes are not neutral. This is usually negligible, as the effects of most synonymous variants are dwarfed by those of disadvantageous nonsynonymous variants, causing the majority of genes to exhibit *d*_N_/*d*_S_ < 1 (Hughes 1999; Holmes 2009). However, this assumption does not hold for overlapping genes (OLGs). A double-stranded nucleic acid may encode up to six protein-coding open reading frames (ORFs), three in the sense direction and three in the antisense direction, allowing pairs of genes to overlap the same nucleotide positions in a genome (Figure 1). In such OLGs, changes that are synonymous in one gene may be nonsynonymous in the other, making otherwise ‘silent’ variants subject to selection. As a result, conventional *d*_N_/*d*_S_ can fail to detect purifying selection or erroneously predict positive (Darwinian) selection when applied to OLGs (Sabath et al. 2008; Sabath and Graur 2010).

OLGs are widespread in viruses (Belshaw et al. 2007; Brandes and Linial 2016; Pavesi et al. 2018), and may not be uncommon in prokaryotes (Meydan et al. 2018; Vanderhaeghen et al. 2018; Weaver et al. 2019) and eukaryotes, including humans (Makałowska et al. 2007; Sanna et al. 2008). The number of OLGs has likely been underestimated, partly because genome annotation software is biased against both short ORFs (Warren et al. 2010) and overlapping ORFs (Vanderhaeghen et al. 2018). Current methods for detecting OLGs, such as Synplot2 (Firth 2014), *d*_N_/*d*_S_ estimators (Sabath et al. 2008; Wei and Zhang 2015), and long-ORF identifiers (Schlub et al. 2018) are subject to one or more of the following limitations: restricted to long OLGs, limited to single or pairs of sequences, unsuitable for low sequence divergence, not specific to protein-coding genes, lacking accessible implementation, or too computationally intensive for genome-scale data. Scalable bioinformatics tools are therefore needed to predict OLG candidates for further analysis, preferably by utilizing the evolutionary information available in multiple sequences and quantifying purifying selection in a way that is comparable to that of non-OLGs. We wrote OLGenie to fill this void.

**Figure 1.**
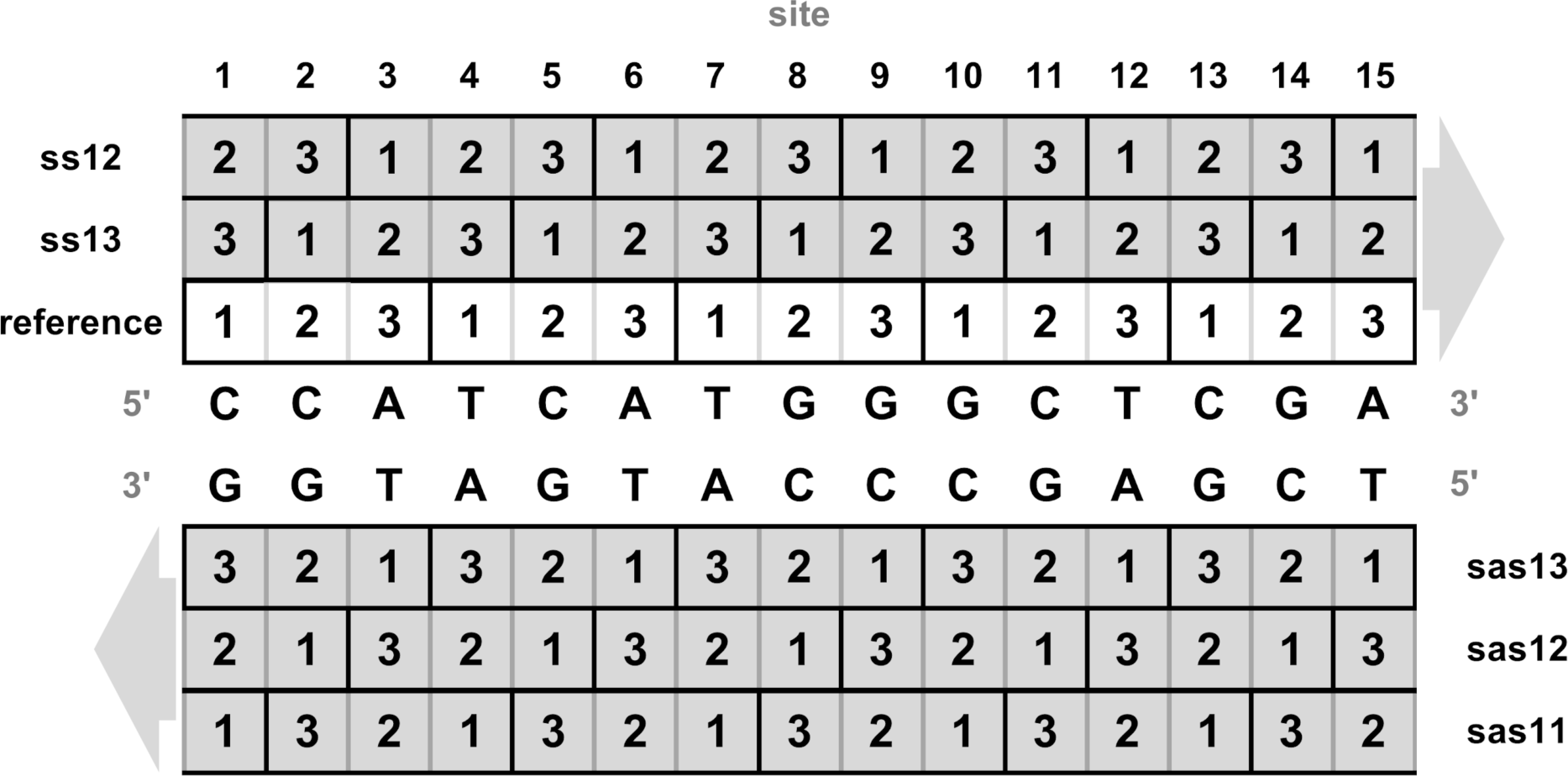
The six possible open (protein-coding) reading frames (ORFs) of a double-stranded nucleic acid sequence. Codons are shown as solid black boxes, each comprising three ordered positions (1, 2, 3). The reference gene frame is shown in white and the five alternate gene frames are shown in grey. To indicate the frame relationship between the reference and alternate genes, we use the nomenclature of Wei and Zhang (2015), where ‘ss’ indicates ‘sense-sense’ (same-strand), ‘sas’ indicates ‘sense-antisense’ (opposite-strand), and the numbers indicate which codon position of the alternate gene (second number) overlaps codon position 1 of the reference gene (first number). Note that the choice of reference frame is arbitrary, and ss12 is equivalent to ss13 except that the reference is swapped, so that no biological differences are expected. Frame sas13 is exceptional in that reference and alternate codons overlap exactly, *i.e.*, one reference codon (positions 1, 2, and 3) overlaps only one alternate codon (positions 3, 2, and 1, respectively). For all other frames, one reference codon partially overlaps each of two alternate codons.

## New Approaches

OLGenie is executed at the Unix/Linux command line with two inputs: (1) a multiple sequence alignment (FASTA file) of contiguous codons known or hypothesized to constitute an OLG pair; and (2) the frame relationship of the OLGs. The codon frame beginning at site 1 of the alignment is considered the ‘reference’ gene, which overlaps an ‘alternate’ gene in one of five frames: ss12, ss13, sas11, sas12, or sas13. Here, ‘ss’ indicates ‘sense-sense’ (same-strand), ‘sas’ indicates ‘sense-antisense’ (opposite-strand), and the numbers indicate which codon position of the alternate gene (second number) overlaps codon position 1 of the reference gene (first number) (Figure 1). The alternate gene is typically shorter, occurring entirely or partially within the reference gene, and is typically of questionable functionality.

OLGenie estimates *d*_N_ and *d*_S_ in OLGs by modifying the method of Wei and Zhang (2015). Four expanded *d*_N_ and *d*_S_ measures are used: *d*_NN_, *d*_SN_, *d*_NS_, and *d*_SS_, where the first subscript refers to the reference gene and the second subscript refers to the alternate gene. For example, *d*_NS_ refers to the number of nucleotide substitutions per site that are nonsynonymous in the reference gene but synonymous in the alternate gene (NS; nonsynonymous-synonymous). Given these values, *d*_N_/*d*_S_ may be estimated for the reference gene as *d*_NN_/*d*_SN_ or *d*_NS_/*d*_SS_, and for the alternate gene as *d*_NN_/*d*_NS_ or *d*_SN_/*d*_SS_. In each case, the effect of mutations in one of the two OLGs is held constant (N or S), ensuring a ‘fair comparison’ in the other gene. For example, if nonsynonymous changes observed in the reference gene are disproportionately synonymous in the alternate gene (*d*_NS_ > *d*_NN_), the result will be *d*_NN_/*d*_NS_ < 1.0, and purifying selection on the alternate gene can be inferred (Hughes and Hughes 2005). In practice, *d*_NN_/*d*_NS_ rather than *d*_SN_/*d*_SS_ is typically used to test for selection in the alternate gene, as SS (synonymous-synonymous) sites are usually too rare to allow a reliable estimate for *d*_SS_.

The original Wei-Zhang method is computationally prohibitive when many nucleotide variants are present in neighboring codons, and the size of the minimal bootstrap unit is data-dependent. To circumvent these issues, we introduce three modifications: (1) consider each reference codon to be an independent unit of the alignment amenable to bootstrapping; (2) apply the Nei-Gojobori method to each OLG, as implemented in SNPGenie (Nei and Gojobori 1986; Nelson et al. 2015; Nelson and Hughes 2015); and (3) consider only observed differences, not all mutational pathways, *i.e.*, two codons either do (synonymous) or do not (nonsynonymous) encode the same amino acid. Modification (1) is not strictly true when two neighboring reference codons share sites with the same alternate codon, introducing biological non-independence. Nevertheless, no individual site is included in more than one unit of the alignment, and the assumption of independence has proven widely effective (Nei and Kumar 2000), even though nearby codons may never evolve independently. Modification (3) is identical to the original Wei-Zhang method when a pair of sequences contains only one difference in contiguous codons. However, synonymous differences may be underestimated when ≥2 sites in contiguous codons differ, as a nonsynonymous change will mask earlier or later synonymous changes. As a result, OLGenie tends to underestimate the denominator of *d*_N_/*d*_S_ (*d*_NS_ or *d*_SN_), biasing the ratio upward and yielding a conservative test of purifying selection that nevertheless has increased power over non-OLG *d*_N_/*d*_S_ (Supplementary Section S1).

## Results and Discussion

### Assessment with Simulated Data

To evaluate OLGenie when selection dynamics are known, we first performed simulation experiments for each frame across a range of *d*_N_/*d*_S_ values, matching sequence divergence to that of our positive controls (median 0.0585; Supplementary Figure S1). Calibration plots reveal that OLGenie produces relatively accurate estimates, especially for purifying selection, with three notable biases: (1) except for sas12, *d*_N_/*d*_S_ is always overestimated; (2) except for sas12, *d*_N_/*d*_S_ overestimation increases when the overlapping gene is under stronger purifying selection; and (3) for sas12, *d*_N_/*d*_S_ is somewhat underestimated for the overlapping gene when *d*_N_/*d*_S_ ≥ 1 (Figure 2; Supplementary Tables S1 and S2). Accuracy improved substantially for lower sequence divergence (Supplementary Figure S2) and suffered minimally at higher transition/transversion ratios (Supplementary Figure S3). Bias (1) is mainly explained by modification (3) above. Bias (2) is explained by the failure to account for unobserved changes (multiple hits), for which no known correction is applicable to OLGs (Hughes et al. 2006), causing the disproportionate underestimation of the denominator (*d*_NS_ or *d*_SN_) in the presence of purifying selection. Bias (3) may be due to the preponderance of “forbidden” codon combinations in sas12, which must necessarily be avoided to prevent STOP codons in the overlapping frame, leading to the overestimation of NN sites and underestimation of *d*_NN_ (Lèbre and Gascuel 2017). Additionally, our observations may be partly attributable to the fact that STOP codons (TAA, TAG, and TGA) are AT-rich and must be avoided in multiple frames to allow OLGs, implicitly favoring high GC content and biasing codon usage (Supplementary Figure S4 and Table S3) (Pavesi et al. 2018). Finally, for all frames, bias and variance are highest when the overlapping gene is under purifying selection.

**Figure 2.**
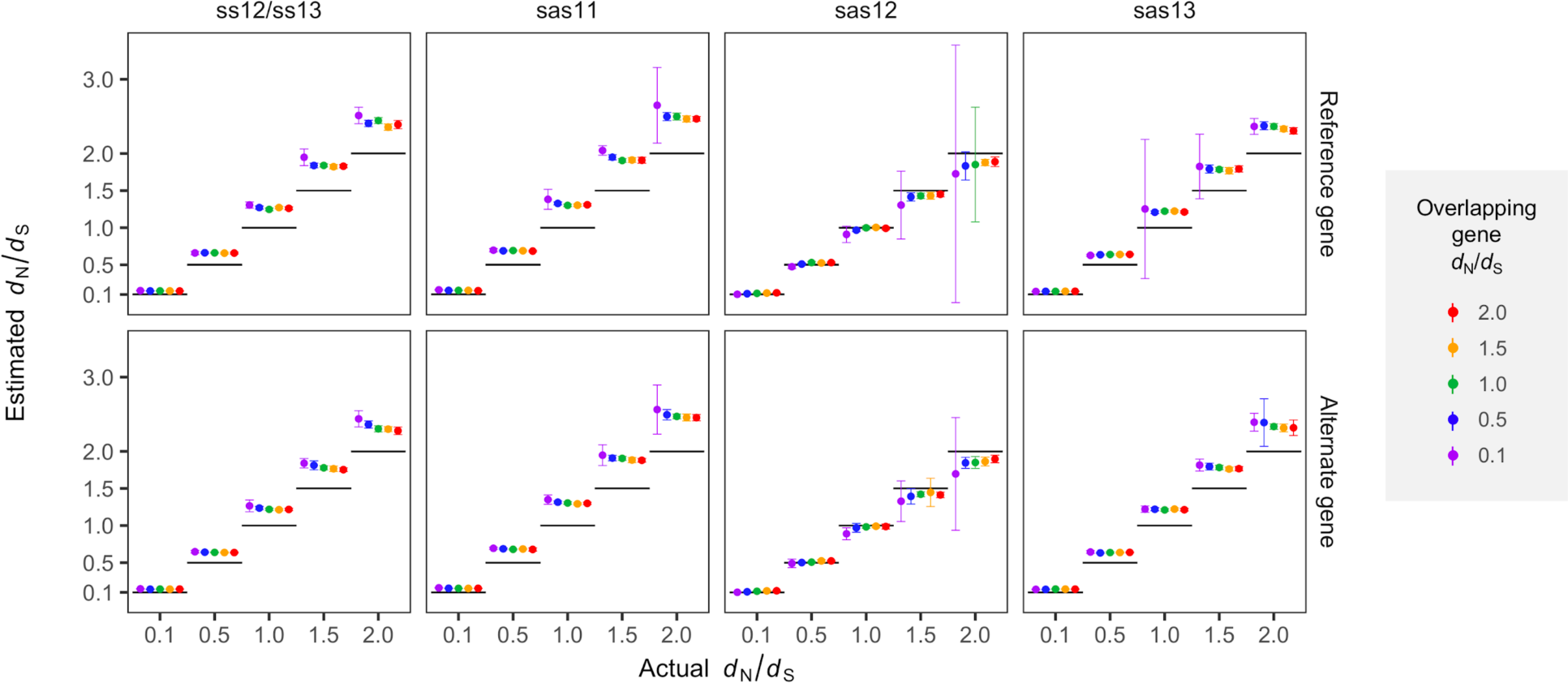
Assessment of OLGenie using simulated sequences. Calibration plots show the accuracy and precision of OLGenie *d*_N_/*d*_S_ estimates for the reference (top row; *d*_NN_/*d*_SN_) and alternate (bottom row; *d*_NN_/*d*_NS_) genes when mean pairwise distance is 0.0585 per site (median of biological controls). For each frame relationship, estimated *d*_N_/*d*_S_ is shown as a function of the actual simulated value, indicated by horizontal black line segments (x axis values), and the *d*_N_/*d*_S_ value of the overlapping gene, indicated by color (left to right: purple = 0.1; blue = 0.5; green = 1.0; orange = 1.5; and red = 2.0). For example, all purple points in the top row refer to simulations with alternate gene *d*_N_/*d*_S_ = 0.1, while all purple points in the bottom row refer to simulations with reference gene *d*_N_/*d*_S_ = 0.1. To obtain highly accurate point estimates, each parameter combination (reference *d*_N_/*d*_S_, alternate *d*_N_/*d*_S_, frame) was simulated using 1,024 sequences of 100,000 codons (Supplementary Table S1). To obtain practical estimates of variance relevant to real OLG data, simulations were again carried out for each parameter combination so as to emulate our biological control dataset: sample size of 234, and sequence length (number of codons) and number of alleles (max 1,024) randomly sampled with replacement from the controls (Supplementary Table S2). Error bars show standard error of the mean, estimated from replicates with defined *d*_N_/*d*_S_ values (≤234). A transition/transversion ratio (*R*) of 0.5 (equal rates) was used; similar results were obtained using *R*=2 (Supplementary Figure S3). Full simulation results are presented in Supplementary Tables S1-S6 and Figures S1-S6.

Our simulations also allowed us to identify the most accurate and precise ratios for estimating each frame’s *d*_N_/*d*_S_. For ss12/ss13, sas11, and sas13, the rarest site class is SS (0-2.7% of sites), leading to high stochastic error when estimating *d*_SS_. Thus, for the alternate gene these frames, the *d*_NN_/*d*_NS_ ratio is relatively ‘site-rich’ and preferred. Contrarily, for sas12, SS sites are usually more common (18.3%) than NS (7.4%) and SN (7.4%) sites, so that *d*_NN_/*d*_NS_ is preferred only 52.5% of the time (51.2 - 53.9%; binomial 95% C.I.) (Supplementary Table S4 and S5; Figures S5 and S6). Thus, for an alternate gene in sas12, either ratio can potentially be informative, and should be selected on a case-by-case basis, based on the number of sites: *d*_NN_/*d*_NS_ if the minimum of (NN, NS) >> minimum of (SN, SS); *d*_SN_/*d*_SS_ if the inequality is reversed; or both if the minima are approximately equal.

### Assessment with Biological Controls

To evaluate OLGenie’s performance with real data, we next applied the program to 58 known OLG (positive control) and 176 non-OLG (negative control) loci from viral genomes (Pavesi et al. 2018). Codon alignments were generated from quality-filtered BLASTN hits (Methods). OLGenie results are 73% accurate (*α* = 0.05), with receiver operating characteristic (ROC) curves yielding an area under the curve (AUC) of 0.70 for the full dataset (Supplementary Table S6). AUC increases only marginally for longer sequences but drastically for lower *d*_N_/*d*_S_ values, reaching AUC = 1.0 for *d*_N_/*d*_S_ ≤ 0.2 (Figure 3; Supplementary Tables S7 and S8). Results are comparable even with less strict alignment criteria (Supplementary Figures S7 and S8 and Tables S9-S12). Importantly, these results may underestimate OLGenie’s performance, as our dataset included more negative than positive controls, and negative controls may include unannotated OLGs. For example, 4 negative controls of length 204-2664nt exhibit *d*_N_/*d*_S_ < 0.2, warranting investigation (Supplementary Table S6). Finally, performance would likely improve for curated alignments limited to carefully defined taxonomic groups.

**Figure 3.**
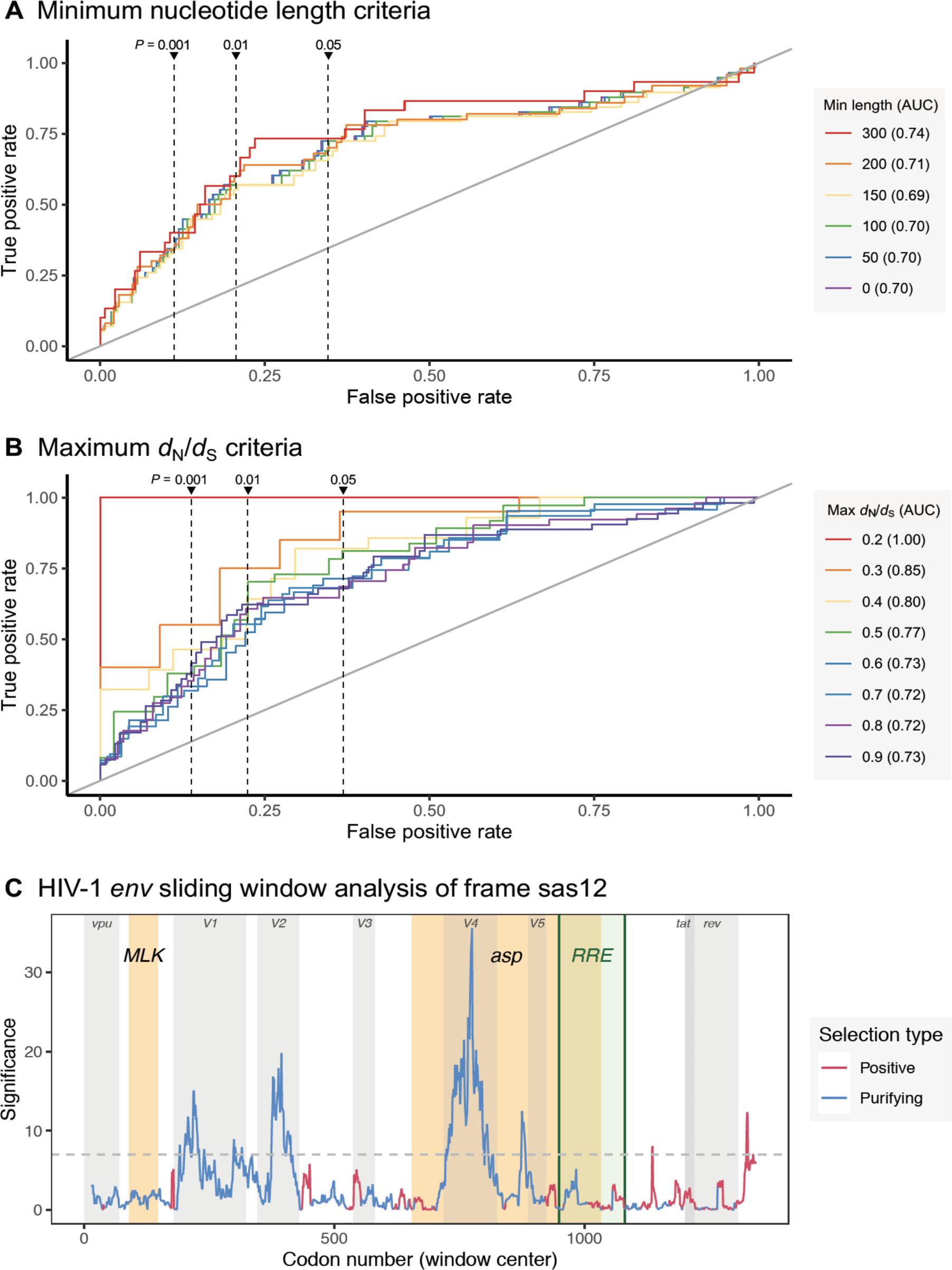

Assessment of OLGenie using biological controls. (*A and B*) Receiver operating characteristic (ROC) curves for overlapping (alternate) gene prediction at varying *P*-value cut-offs (y axis: true positive rate=sensitivity; x axis: false positive rate=[1 - specificity]). Curves were plotted for subsets of the data corresponding to differing minimum length (*A*) and maximum *d*_N_/*d*_S_ (*B*) criteria, following the approach of Schlub et al. (2018), with red indicating the strictest criteria. The full dataset is represented by purple in (*A*) (overlaps blue), with area under the curve (AUC) reported in parentheses in the key (Supplementary Tables S6-S8). The ROC expected using random classification (AUC = 0.5) is shown as a diagonal grey line. Vertical dashed lines show mean false positive rates for *P*-value cut-offs of 0.001, 0.01, and 0.05 (left to right). The site-rich *d*_NN_/*d*_NS_ ratio was used to analyze 234 controls (81 ss12 and 153 ss13): 58 positive (16 ss12 and 42 ss13) and 176 negative (65 ss12 and 111 ss13). Of these, 162 (30 positive, 132 negative) had length ≥300nt and 14 (10 positive, 4 negative) had *d*_N_/*d*_S_ ≤0.2. (*C*) The HIV-1 *env* gene was analyzed in sas12 with the site-rich ratio *d*_NN_/*d*_NS_ using 25-codon sliding windows (step size=1). Significance (y axis) shows the natural logarithm of the inverse *P*-value, calculated using *Z*-tests (1,000 bootstrap replicates per window). The horizontal dashed grey line shows the multiple comparisons *P*-value threshold (0.000924) described in Supplementary Section S5. Results for other frames are shown in Supplementary Figure S9. Positive selection (red) refers to *d*_N_/*d*_S_ > 1; purifying selection (blue) refers to *d*_N_/*d*_S_ < 1. Sequence features are described in Supplementary Table S15 and shown here as shaded rectangles: yellow for hypothesized sas12 genes, green for the highly structured RNA Rev response element (RRE), and grey otherwise.

### Case Study: HIV-1’s Putative *antisense protein* Gene

Lastly, we examined the unresolved case of HIV-1’s *env*/*asp* sas12 overlap (Miller 1988; Torresilla et al. 2015), where the putative *antisense protein* (*asp*) gene has evaded detection by several methods, including non-OLG *d*_N_/*d*_S_ (Cassan et al. 2016; Schlub et al. 2018). We used OLGenie to test for purifying selection in three subregions of *env*: (1) 5′ non-OLG; (2) putative *asp*-encoding; and (3) 3′ non-OLG. Three datasets were used: (1) M group from Cassan et al. (2016) (1,723 codons × 23,831 sequences; functional *asp* hypothesized); (2) non-M groups from Cassan et al. (1,723 codons × 92 sequences; no functional *asp* hypothesized); and (3) HIV-1 BLAST hits for *env* using the same methods as our control dataset (1,355 codons × 4,646 sequences). We employed *d*_NN_/*d*_NS_ for the alternate gene, as this ratio was far and away the most site-rich for all *env* frames (*e.g.*, sas12 site counts: NN=2,127.2 and NS=825.3, *versus* SN=190.1 and SS=636.4; Supplementary Table S13).

The sas12 *d*_N_/*d*_S_ ratio was significantly smaller than 1 in all three datasets for the 5′ non-OLG (*d*_N_/*d*_S_ ≤ 0.66; *P* = 2.04 × 10^−7^) and *asp* (*d*_N_/*d*_S_ ≤ 0.58; *P* = 2.75 × 10^−5^) subregions of *env*. The lowest ratio for each dataset always occurred in *asp*, reaching very high significance in the BLAST dataset (*d*_N_/*d*_S_ = 0.29; *P* = 5.04 × 10^−25^). As a comparison, our ss12/ss13 controls suggest a false positive rate of 0% for *d*_N_/*d*_S_ ≤ 0.4 when employing *P* ≤ 1.04 × 10^−6^ (based on 28 OLGs and 27 non-OLGs). The 3′ non-OLG region was also significant for the Cassan non-M groups (*d*_N_/*d*_S_= 0.78, *P* = 0.00921); however, the expected false positive rate is ~22-28% and the other two datasets are not significant (*d*_N_/*d*_S_ ≥ 0.74; *P* ≥ 0.107) (Supplementary Table S14).

To test whether our results were an artifact of other sequence features, including the highly structured RNA Rev response element (RRE; Supplementary Table S15; Fernandes et al. 2012), we also used OLGenie to perform sliding window analyses. Results show that purifying selection in *env*/sas12 is most significant in regions of *asp* not overlapping the RRE (Figure 3C). The strongest evidence is observed in V4, suggesting that nonsynonymous changes in this region are disproportionately synonymous in *asp*. Significance is also attained for the two known ss12 OLGs, *vpu* and *rev* (Supplementary Figures S9 and S10). Thus, OLGenie specifically detects protein-coding function in all three datasets. Contrarily, Synplot2 shows the strongest evidence for synonymous constraint in the RRE, likely due to RNA structure rather than protein-coding function, and fails to detect *vpu* in the BLAST dataset (Supplementary Figure S11). These results suggest that purifying selection acts on the sas12 protein-coding frame of *env*, particularly in the *asp* region.

## Conclusions

OLGenie provides a simple, accessible, scalable method for estimating *d*_N_/*d*_S_ in OLGs. It utilizes a well-understood measure of natural selection that is specific to protein-coding genes, making it possible to directly compare functional constraint between OLGs and non-OLGs. Moreover, although its estimates of constraint are conservative, its discriminatory ability exceeds that of other methods (*e.g.*, Schlub et al. 2018). Power is greatest at relatively low levels of sequence divergence, and may be increased in the future by incorporating mutational pathways or comparing conservative *vs.* radical nonsynonymous changes. Even so, not all functional genes exhibit detectable selection, so that some OLGs are likely to be missed by any selection-based method. Nevertheless, because candidate OLGs are usually subject to costly downstream laboratory analyses, minimizing the false positive rate is paramount. To this end, OLGenie achieves a false positive rate of 0% for several subsets of our control data, *e.g.*, regions with *d*_N_/*d*_S_ < 0.4 and *P* ≤ 1.04 × 10^−6^. OLGenie can therefore be used to predict OLG candidates with high confidence, allowing researchers to begin studying evolutionary evidence for OLGs at the genomic scale.

## Methods

OLGenie is written in Perl with no dependencies, and is freely available at https://github.com/chasewnelson/OLGenie. Estimates of *d* are obtained by calculating *d*_NN_ = *m*_NN_/*L*_NN_, *d*_SN_ = *m*_SN_/*L*_SN_, *d*_NS_ = *m*_NS_/*L*_NS_, or *d*_SS_ = *m*_SS_/*L*_SS_, where *m* is the mean number of differences and *L* is the mean number of sites between all allele pairs at each reference codon. Simulation scripts were modified from Wei and Zhang (2015). Biological control gene coordinates were obtained from Pavesi et al. (2018) and used to retrieve nucleotide sequences from the latest NCBI genome. Homologous sequences were obtained using BLASTN (Altschul et al. 1990); excluded if they contained in-frame STOP codons or were <70% of query length (Hughes et al. 2005); translated; aligned using MAFFT v.7.150b (Katoh and Standley 2013); codon-aligned using PAL2NAL v14 (Suyama et al. 2006); and filtered to keep only unique alleles and codon positions with ≥6 defined (non-gap) sequences (Jordan and Goldman 2012). Statistical analyses were carried out in R v3.5.2 (R Core Team 2018). Significant deviations from *d*_N_ *-d*_S_ = 0 were detected using *Z*-tests for genes and sliding windows (10,000 and 1,000 bootstrap replicates, respectively). Complete methods, results, and data are available in the Supplementary Material.

## Author Contributions

C.W.N. conceived and wrote OLGenie, obtained control data, processed data, performed OLGenie analyses, created figures, modified the simulation software, and drafted the manuscript. Z.A. obtained and processed HIV-1 example data and performed Synplot2 analyses. X.W. wrote the simulation software and advised on statistical and sequence analyses. C.W.N. and X.W. developed statistical methods. All authors designed the study, analyzed and interpreted data, and revised the manuscript.

## Supplementary Material

Supplementary material is available online at https://doi.org/10.5281/zenodo.3575391.

## Acknowledgments and Funding

This study was supported by a Gerstner Scholars Fellowship from the Gerstner Family Foundation at the American Museum of Natural History to C.W.N., and funding from the Bavarian State Government and National Philanthropic Trust to Z.A. The authors thank Wen-Hsiung Li for comments on an earlier draft of this manuscript; Ming-Hsueh Lin for dramatic improvements to our visuals; and Reed A. Cartwright, Robert S. Harbert, Jim Hussey, Michael Lynch, Sergios Orestis-Kolokotronis, Apurva Narechania, Sally Warring, Jeff Witmer, Meredith Yeager, Jianzhi (George) Zhang, Martine Zilversmit, and the Sackler Institute for Comparative Genomics workgroup for discussion.

## Notes

https://github.com/chasewnelson/OLGenie

https://doi.org/10.5281/zenodo.3575391

